# Testing structural models of psychopathology at the genomic level

**DOI:** 10.1101/502039

**Authors:** Irwin D. Waldman, Holly E. Poore, Justin M. Luningham, Jingjing Yang

## Abstract

Genome-wide association studies (GWAS) have revealed hundreds of genetic loci associated with the vulnerability to major psychiatric disorders, and post-GWAS analyses have shown substantial genetic correlations among these disorders. This evidence supports the existence of a higher-order structure of psychopathology at both the genetic and phenotypic levels. Despite recent efforts by collaborative consortia such as the Hierarchical Taxonomy of Psychopathology (HiTOP), this structure remains unclear. In this study, we tested multiple alternative structural models of psychopathology at the genomic level, using the genetic correlations among fourteen psychiatric disorders and related psychological traits estimated from GWAS summary statistics. The best-fitting model included four correlated higher-order factors – externalizing, internalizing, thought problems, and neurodevelopmental disorders – which showed distinct patterns of genetic correlations with external validity variables and accounted for substantial genetic variance in their constituent disorders. A bifactor model including a general factor of psychopathology as well as the four specific factors fit worse than the above model. Several model modifications were tested to explore the placement of some disorders – such as bipolar disorder, obsessive-compulsive disorder, and eating disorders – within the broader psychopathology structure. The best-fitting model indicated that eating disorders and obsessive-compulsive disorder, on the one hand, and bipolar disorder and schizophrenia, on the other, load together on the same thought problems factor. These findings provide support for several of the HiTOP higher-order dimensions and suggest a similar structure of psychopathology at the genomic and phenotypic levels.

---

Over the past several years, genome-wide association studies (GWAS) have shed considerable light on the genetic underpinnings of major psychiatric disorders, including schizophrenia, bipolar disorder, and depression^1–3^. In addition to revealing replicable genetic loci associated with these disorders, various post-GWAS analyses have identified the amount of trait variation that is due to genetic factors – i.e., the single nucleotide polymorphism (SNP)-based heritability^4,5^ – as well as the genetic correlations between traits^6^. Recent studies have shown substantial genetic correlations among various psychiatric disorders^6,7^, mirroring phenotypic correlations, and suggesting a shared genetic vulnerability which reflects a higher-order structure of psychopathology^8–10^.

Various models of the underlying phenotypic structure of psychopathology, which capture the substantial correlations among psychiatric disorders, have been advanced in the literature, including a two-factor model comprising externalizing and internalizing dimensions^11^, a three factor model that distinguishes distress from fears within internalizing^12^, and models that include a thought problems factor^13^.

One theoretical conceptualization of the structure of psychopathology, the Hierarchical Taxonomy of Psychopathology (HiTOP)^8^, posits that the risk for psychopathology is captured by a general factor (p factor), which in turn influences specific spectra of psychopathology (e.g., internalizing, thought disorder), which in turn influence more specific dimensions or subfactors (e.g., fears and distress pathology) and disorders (e.g., major depressive disorder).

A bifactor model, including a general factor onto which all disorders load and specific factors that capture the remaining covariance related to groups of disorders (e.g., externalizing and internalizing), has shown a sharp rise in popularity among psychopathology researchers. Nonetheless, statisticians have pointed out difficulties in distinguishing between bifactor and correlated factor models^14,15^ and the tendency for model fit indices to be biased in favor of the bifactor model^16–18^.

Some researchers argue that genetic and psychobiological levels of analysis enhance investigation of the structure of psychopathology and augment what is learned through pure statistical comparisons^14^. Given this, and the moderate-to-high genetic correlations observed among psychiatric disorders and related psychological traits, examination of the higher-order structure of psychopathology at the genomic level is warranted.

Two recent studies have examined the factor structure of psychopathology and related traits at the genomic level. Grotzinger et al^19^ fit a model containing a single common factor of psychopathology using GWAS summary statistics for schizophrenia, bipolar disorder, major depressive disorder (MDD), post-traumatic stress disorder (PTSD), and anxiety disorders. Their results indicated that each disorder had a moderate-to-high loading on the common factor, revealing that genetic covariation among psychiatric disorders can be captured using factor analysis. Lee et al^20^ used an exploratory approach to examine the genetic covariance among eight psychiatric disorders using GWAS summary statistics, and found evidence for a three-factor model which included factors representing compulsive behaviors, mood and psychotic disorders, and neurodevelopmental disorders.

In the present study, we capitalized on the fourteen largest GWAS of psychiatric disorders and related psychological traits to obtain estimates of genetic correlations and test alternative structural models of psychopathology at the genomic level. We included more disorders and traits and tested more alternative models of psychopathology than in previous studies^19,20^, guided by both the phenotypic literature and previously estimated genetic correlations. We also evaluated the construct validity of our best-fitting model by estimating genetic correlations between the higher-order factors and external criterion variables, such as educational attainment and personality characteristics.

## METHODS

### GWAS summary statistics

We conducted a systematic search of repositories of publicly available GWAS summary statistics for psychiatric disorders and relevant external criterion variables.

The summary statistics for attention-deficit/hyperactivity disorder (ADHD)^21^, autism spectrum disorder (ASD)^22^, bipolar disorder^3^, anorexia nervosa^23^, MDD^1^, schizophrenia^2^, PTSD^24^, obsessive-compulsive disorder (OCD)^25^, tobacco use^26^, and anxiety disorders^27^ were downloaded from the Psychiatric Genomics Consortium (PGC) repository. Some of these samples were augmented by samples from other consortia, such as the Lundbeck Foundation Initiative for Integrative Psychiatric Research (iPSYCH) for ADHD, ASD and MDD; the Anxiety NeuroGenetics Study (ANGST) for anxiety disorders; the International Obsessive Compulsive Disorder Foundation Genetics Collaborative (IOCDF-GC) and OCD Collaborative Genetics Association Studies (OCGAS) for OCD; and the Tobacco and Genetics Consortium (TGC) for tobacco use. The summary statistics for antisocial behavior^28^ were obtained from the Broad Antisocial Behavior Consortium (BroadABC), and those for aggression^29^ from the Early Genetics and Lifecourse Epidemiology (EAGLE) consortium (Table 1).

**Table 1.** Characteristics of studies of disorders and traits included in the analyses

| Phenotype | Study | Year | Data source | Sample size<br>(cases/controls) | Study design | Effect<br>size | Ancestry |
| --- | --- | --- | --- | --- | --- | --- | --- |
| ADHD | DeMontis et al<sup>21</sup> | 2017 | PGC, iPSYCH | 20,183 / 35,191 | Case-control | OR | European |
| Alcohol dependence | Walters et al<sup>37</sup> | 2018 | PGC | 10,206 / 28,480 | Case-control | OR | European |
| Cannabis<br>dependence | Agrawal et al<sup>38</sup> | 2018 | PGC | 3,757 / 9,931 | Case-control | Beta | European |
| Tobacco use | Furberg et al<sup>26</sup> | 2010 | PGC, TGC | 73,853 | Continuous | Beta | European |
| Aggression | Pappa et al<sup>30</sup> | 2016 | EAGLE | 18,988 | Continuous | Beta | European |
| Antisocial behavior | Tielbeek et al<sup>28</sup> | 2017 | BroadABC | 16,400 | Continuous | Beta | European |
| Eating disorders | Duncan et al<sup>24</sup> | 2017 | PGC | 3,495 / 10,982 | Case-control | OR | European |
| Anxiety disorders | Otowa et al<sup>27</sup> | 2016 | PGC, ANGST | 7,016 / 14,745 | Case-control | Beta | European |
| PTSD | Duncan et al<sup>24</sup> | 2017 | PGC | 2,424 / 7,113 | Case-control | OR | European |
| MDD | Wray et al<sup>1</sup> | 2018 | PGC, iPSYCH | 59,851 / 113,154 | Case-control | OR | European |
| OCD | Arnold et al<sup>25</sup> | 2017 | PGC, IOCDF-GC,<br>OCGAS | 2,688 / 7,037 | Case-control | OR | European |
| Schizophrenia | Ripke et al<sup>2</sup> | 2014 | PGC | 36,989 / 113,075 | Case-control | OR | Admixed |
| Bipolar disorder | Stahl et al<sup>3</sup> | 2019 | PGC | 20,352 / 31,358 | Case-control | OR | European |
| ASD | Grove et al<sup>22</sup> | 2019 | PGC, iPSYCH | 18,382 / 27,969 | Case-control | OR | European |

ADHD – attention-deficit/hyperactivity disorder, PTSD – post-traumatic stress disorder, MDD – major depressive disorder, OCD – obsessive-compulsive disorder, ASD – autism spectrum disorder, PGC – Psychiatric Genetics Consortium, iPsych – Lundbeck Foundation Initiative for Integrative Psychiatric Research, EAGLE – Early Genetics and Lifecourse Epidemiology, BroadABC – Broad Antisocial Behavior Consortium, ANGST – Anxiety NeuroGenetics Study, IOCDF-GC – International Obsessive Compulsive Disorder Foundation Genetics Collaborative, OCGAS – OCD Collaborative Genetics Association Studies, TGC – Tobacco and Genetics Consortium, OR – odds ratio, Beta – standardized regression coefficient

The summary statistics for age at first birth and number of children^30^, neuroticism, subjective well-being, depression symptoms^31^, and educational attainment^32^ were downloaded from the Social Science Genetic Association Consortium (SSGAC) repository; those for extraversion^33^, openness to experience, agreeableness, and conscientiousness^34^ from the Genetics of Personality Consortium (GPC) repository; those for loneliness^35^ from the PGC; and those for body mass index^36^ were obtained from the Genetic Investigation of Anthropometric Traits (GIANT) consortium repository and the UK Biobank (Table 2).

**Table 2.** Characteristics of external criterion variables

| Phenotype | Study | Year | Data source | Sample size | Study design | Effect size | Ancestry |
| --- | --- | --- | --- | --- | --- | --- | --- |
| Neuroticism | Okbay et al<sup>31</sup> | 2016 | SSGAC | 298,420 | Continuous | Beta | European |
| Depression symptoms | Okbay et al<sup>31</sup> | 2016 | SSGAC | 161,460 | Continuous | Beta | European |
| Subjective well-being | Okbay et al<sup>31</sup> | 2016 | SSGAC | 298,420 | Continuous | Beta | European |
| Extraversion | Van Den Berg et al<sup>33</sup> | 2015 | GPC | 170,910 | Continuous | Beta | European |
| Agreeableness | De Moor et al<sup>34</sup> | 2012 | GPC | 20,669 | Continuous | Beta | European |
| Conscientiousness | De Moor et al<sup>34</sup> | 2012 | GPC | 20,669 | Continuous | Beta | European |
| Openness | De Moor et al<sup>34</sup> | 2012 | GPC | 20,669 | Continuous | Beta | European |
| Educational attainment | Lee et al<sup>32</sup> | 2018 | SSGAC | 766,345 | Continuous | Beta | European |
| Loneliness | Gao et al<sup>35</sup> | 2016 | PGC | 10,760 | Continuous | Beta | Admixed |
| Body mass index | Yengo et al<sup>36</sup> | 2018 | GIANT + UK<br>Biobank | 681,275 | Continuous | Beta | European |
| Number of children | Barban et al<sup>30</sup> | 2016 | SSGAC | 333,702 | Continuous | Beta | European |
| Age at first birth | Barban et al<sup>30</sup> | 2016 | SSGAC | 237,516 | Continuous | Beta | European |

SSGAC – Social Science Genetic Association Consortium, GPC – Genetics of Personality Consortium, PGC – Psychiatric Genetics Consortium, GIANT – Genetic Investigation of Anthropometric Traits consortium, Beta – standardized regression coefficient

When summary statistics for an existing GWAS could not be found online, the authors of the relevant publications were contacted via email and asked to provide summary statistics, as was the case for alcohol dependence^37^ and cannabis dependence^38^. When results from more than one GWAS of the same disorder were available, the most recent and largest GWAS was chosen. With the exception of schizophrenia and loneliness, for which only GWAS from admixed populations were available, we used summary statistics from European ancestry individuals.

Tobacco use, antisocial behavior, aggression, and all of the external criterion variables were assessed using a continuous variable study design, whereas GWAS for all other psychiatric disorders used a case-control design. The total sample size included in analyses consisted of 658,640 participants.

### Statistical analysis

All analyses were conducted using the recently developed Genomic Structural Equation Modeling (SEM) R package^19^. Genomic SEM employs a novel extension of the widely used LD-score regression method^4^ that calculates the genetic covariance among traits using GWAS summary statistics from multiple studies. Potential sample overlap across studies (e.g., shared control samples) is accounted for by the regression.

Genomic SEM first estimates a *p x p* genetic covariance matrix S containing SNP-based heritabilities for each of the *p* disorders or traits on the diagonal and genetic covariances among the *p* disorders and traits in the off-diagonal elements. The estimation uncertainty of S that is required for accurate model estimation is captured in a matrix V, which contains squared standard errors of the estimates in S on the diagonal, and the covariance between each pair of elements of S in the off-diagonal. These off-diagonal terms capture the potential sample overlap across traits.

After GWAS summary statistics were identified for the fourteen psychiatric disorders and traits of interest, the publicly-available files were formatted for Genomic SEM pre-processing. Next, the genetic covariance matrix was calculated using LD weights for populations of European ethnicity provided by the Broad Institute and the LD-score regression function of the Genomic SEM R package. The estimated genetic covariance matrix S and its associated sampling matrix V were then used for model fitting analyses.

Pre-specified structural models were fitted and evaluated using the weighted least squares (WLS) discrepancy function. WLS directly incorporates the V matrix, and is also recommended over maximum likelihood estimation by the creators of Genomic SEM^19^.

The alternative *a priori* hypothesized structural models were fitted and compared utilizing confirmatory factor analysis (CFA). To evaluate the fit of each model, we used the comparative fit index (CFI) and the Bayesian information criterion (BIC). The fit of each model was evaluated using the combination of CFI and BIC, as each individual fit index has its strengths and limitations and a consensus has not been reached to use a single index to evaluate the adequacy of model fit^39^.

The CFI is an absolute index of model fit where values >0.90 indicate good fit^40,41^, whereas the BIC is a relative index of model fit that can be used to adjudicate among alternative models^39,42^. The model with the lowest value for BIC is considered the best fitting model, and it has been shown that differences of BIC >10 represent very strong evidence in favor of the model with the lower BIC^43^. Models were considered untenable if they contained factor loadings that were out of bounds, not significantly different from zero, or had very large standard errors.

User-defined models were provided to the genomic SEM software in the lavaan syntax^44^. Six CFA models with increasing complexity were specified *a priori* to evaluate and contrast different hypotheses regarding the latent factor structure of psychopathology. The alternative models were defined as specified below.

Model 1 included a single common factor on which all disorders and traits loaded. Model 2 was characterized by three correlated psychopathology factors (externalizing, internalizing, and thought problems). Externalizing was indicated by ADHD, aggression, alcohol dependence, cannabis dependence, tobacco use, and antisocial behavior; internalizing was indicated by MDD, PTSD, anxiety disorders, and eating disorders; and thought problems was indicated by schizophrenia, bipolar disorder, OCD and ASD.

Model 3 included four correlated factors representing externalizing, internalizing, thought problems, and substance use disorders. Model 4 posited a four-factor structure extending Model 2, in which neurodevelopmental disorders – i.e., ADHD, ASD, and aggression – loaded onto a unique factor. In this model, the neurodevelopmental disorders were specified with what are known as cross-loadings: they were indicators of the same factors from the previous three-factor model as well as of the new unique factor, which was uncorrelated with the other three factors. Model 5 was similar to model 4, except that ADHD, ASD and aggression loaded only on the neurodevelopmental disorders factor, which was correlated with all the other factors.

Model 6 specified a bifactor model with a general psychopathology factor and four uncorrelated specific factors (externalizing, internalizing, thought problems, and neurodevelopmental disorders). In this model, all disorders loaded on a general factor as well as on their respective specific factors, which were orthogonal to the general factor and to each other. This structure implies that the correlations among all disorders and traits across psychopathology domains are only due to the general factor, whereas the correlations among disorders and traits within psychopathology domains are also due to the domain-specific factors.

Several exploratory models were also tested (Models 5a-5h), due to conflicting evidence in the literature regarding the placement of individual disorders (bipolar disorder, OCD, MDD and eating disorders) within the larger multivariate psychopathology structure. All exploratory models were tested as variations of Model 5.

Finally, we estimated genetic correlations of the higher-order psychopathology factors with several external criterion variables. These genetic correlations were estimated within the measurement model such that disorders’ loadings on their respective factors as well as the higher-order factors’ correlations with external criterion variables were simultaneously estimated in Genomic SEM.

## RESULTS

Genetic correlations among the fourteen psychiatric disorders and related traits are shown in Table 3. Correlations among disorders are strongest within each psychopathology domain (externalizing, internalizing, thought problems, and neurodevelopmental disorders). However, correlations among disorders across psychopathology domains are non-negligible and in some cases of moderate magnitude.

**Table 3.** Genetic correlations among the fourteen psychiatric disorders and related traits

|  | AGG | ADHD | ASD | CIGS | CAN | ALC | ASB | ANX | MDD | PTSD | BIP | OCD | SCZ | ED |
| --- | --- | --- | --- | --- | --- | --- | --- | --- | --- | --- | --- | --- | --- | --- |
| AGG | 1 |  |  |  |  |  |  |  |  |  |  |  |  |  |
| ADHD | 0.77 | 1 |  |  |  |  |  |  |  |  |  |  |  |  |
| ASD | 0.49 | 0.37 | 1 |  |  |  |  |  |  |  |  |  |  |  |
| CIGS | 0.52 | 0.41 | 0.07 | 1 |  |  |  |  |  |  |  |  |  |  |
| CAN | 0.81 | 0.42 | 0.03 | 0.12 | 1 |  |  |  |  |  |  |  |  |  |
| ALC | 0.12 | 0.41 | 0.02 | 0.33 | 0.12 | 1 |  |  |  |  |  |  |  |  |
| ASB | 0.24 | 0.52 | 0.21 | 0.20 | 0.41 | 0.59 | 1 |  |  |  |  |  |  |  |
| ANX | 0.67 | 0.30 | 0.28 | 0.09 | 0.35 | 0.54 | 0.42 | 1 |  |  |  |  |  |  |
| MDD | 0.46 | 0.56 | 0.44 | 0.16 | 0.23 | 0.44 | 0.55 | 0.89 | 1 |  |  |  |  |  |
| PTSD | 0.40 | 0.52 | 0.24 | 0.44 | 0.41 | 0.33 | 0.36 | 0.09 | 0.49 | 1 |  |  |  |  |
| BIP | 0.11 | 0.12 | 0.13 | 0.20 | 0.34 | 0.24 | 0.11 | 0.18 | 0.33 | 0.07 | 1 |  |  |  |
| OCD | 0.38 | -0.16 | 0.12 | -0.05 | 0.25 | -0.27 | -0.05 | 0.30 | 0.30 | 0.42 | 0.32 | 1 |  |  |
| SCZ | 0.04 | 0.12 | 0.21 | 0.11 | 0.07 | 0.09 | 0.09 | 0.26 | 0.37 | 0.18 | 0.68 | 0.33 | 1 |  |
| ED | -0.20 | -0.26 | -0.08 | -0.12 | 0.04 | -0.10 | -0.10 | 0.09 | 0.20 | -0.02 | 0.18 | 0.50 | 0.23 | 1 |

AGG – aggression, ADHD – attention-deficit/hyperactivity disorder, ASD – autism spectrum disorder, CIGS – number of cigarettes smoked per day, CAN – cannabis dependence, ALC – alcohol dependence, ASB – antisocial behavior, ANX – anxiety disorders, MDD – major depressive disorder, PTSD – post-traumatic stress disorders, BIP – bipolar disorder, OCD – obsessive-compulsive disorder, SCZ – schizophrenia, ED – eating disorders. Borders denote correlations among disorders within each higher-order dimension

Fit statistics of the alternative models reflecting the underlying structure of psychopathology are presented in Table 4. We first contrasted a model with a single common factor (Model 1) with a three-factor model that comprised externalizing, internalizing, and thought problems dimensions (Model 2). Model 2 had significantly better fit than Model 1, based on a CFI closer to 0.90 and a much smaller BIC. Next we tested Models 3, 4 and 5, all of which included four factors. Model 3 (including externalizing, internalizing, thought problems, and substance use disorders) resulted in a larger BIC value than Model 2, indicating that the addition of the substance use disorders factor resulted in a worse-fitting model. In contrast, Model 4 (specifying a neurodevelopmental disorders factor uncorrelated with externalizing, internalizing, and thought problems factors) fit better than the three correlated factor model, based on a large reduction in BIC. Model 5, in which the neurodevelopmental disorders factor was correlated with the other factors, resulted in a CFI above 0.90 and another substantial reduction in BIC. Finally, Model 6 (a bifactor model that comprised a general factor as well the four specific externalizing, internalizing, thought problems, and neurodevelopmental disorders factors, all of which were uncorrelated) fit worse than Model 5.

**Table 4.** Models and model fit statistics

| | $\chi^2$ | df | CFI | BIC | Model<br>compared to | ΔBIC |
| --- | --- | --- | --- | --- | --- | --- |
| 1. One common factor | 1052 | 77 | 0.71 | 1427.0 |  |  |
| 2. Three correlated factors (EXT, INT and TP) | 554 | 74 | 0.86 | 969.3 | 1 | 457.7 |
| 3. Four correlated factors (EXT, INT, TP and SUD) | 548 | 71 | 0.86 | 1003.5 | 2 | +34.2 |
| 4. Four factor model (EXT, INT, TP and uncorrelated NDD) | 419 | 71 | 0.90 | 874.5 | 2 | 94.8 |
| 5. Four correlated factors (EXT, INT, TP and NDD; ED on INT) | 385 | 71 | 0.91 | 840.5 | 4 | 34.0 |
| 6. Bifactor model, with four uncorrelated specific factors<br>(ED on INT only) | 400 | 63 | 0.90 | 962.7 | 5 | +122.2 |
| Modified four correlated factors models |  |  |  |  |  |  |
| 5a. Four correlated factors, BIP on TP and EXT | 384 | 70 | 0.91 | 852.9 | 5 | 12.4 |
| 5b. Four correlated factors, BIP on TP and INT | 383 | 70 | 0.91 | 851.9 | 5 | 11.4 |
| 5c. Four correlated factors, OCD on TP and INT | 385 | 70 | 0.91 | 853.9 | 5 | 13.4 |
| 5d. Four correlated factors, OCD on INT only | 402 | 71 | 0.90 | 857.5 | 5 | 17.0 |
| 5e. Four correlated factors, MDD on INT and TP | 382 | 70 | 0.91 | 850.9 | 5 | 10.4 |
| 5f. Four correlated factors, ED on INT and EXT | model would not run |  |  |  |  |  |
| 5g. Four correlated factors, ED on INT and TP | 341 | 70 | 0.92 | 809.9 | 5 | 30.6 |
| **5h. Four correlated factors, ED on TP only** | **341** | **71** | **0.92** | **796.5** | **5** | **44.0** |
| 5i. Bifactor model, with four uncorrelated specific factors<br>(ED on TP only) | 366 | 63 | 0.91 | 928.7 | 5h | +132.2 |

CFI – comparative fit index, BIC – Bayesian information criterion, EXT – externalizing factor, INT – internalizing factor, TP – thought problems factor, SUD – substance use disorders factor, NDD – neurodevelopmental disorders factor, BIP – bipolar disorder, OCD – obsessive-compulsive disorder, MDD – major depressive disorder, ED – eating disorders. The model in bold is the best-fitting model based on a BIC difference >10. $\Delta$ BICs that have plus signs indicate that the more parsimonious models have better fit than the more complex models.

Models in which bipolar disorder loaded on thought problems and externalizing (Model 5a) or thought problems and internalizing (Model 5b) were rejected, as they fit worse than Model 5, and due to bipolar disorder’ s small and negative factor loadings on the externalizing and internalizing factors (i.e., –.01, SE=.10 and –.05, SE=.10, respectively).

A model in which OCD loaded on thought problems and internalizing (Model 5c) was rejected because it fit worse than Model 5 and due to OCD’s small and non-significant loading on internalizing (i.e., .06, SE=.09). Similarly, a model in which OCD loaded only on internalizing (Model 5d) fit worse than a model in which it loaded only on thought problems.

A model in which MDD loaded on internalizing and thought problems (Model 5e) was rejected because it fit worse than Model 5 and because MDD had a negative loading on thought problems (–.10, SE=.15) and its loading on internalizing was out of bounds (1.05, SE=.17).

Models in which eating disorders loaded on internalizing and externalizing (Model 5f) or internalizing and thought problems (Model 5g) were rejected either because they would not run (Model 5f) or due to a negative loading on internalizing (Model 5g: –.27, SE=.08). However, a model in which eating disorders loaded only on thought problems (Model 5h) had a better fit than Model 5, and eating disorders loaded most strongly in this model compared to any other model tested.

We also tested a bifactor version of this model (Model 5i), which fit worse than Model 5 and had problematic model characteristics. Specifically, as shown in Table 5, all of the disorders’ loadings on the externalizing and internalizing specific factors became non-significant and some loadings became negative (cannabis and PTSD) after accounting for their loading on the general factor, while the loading of eating disorders on the general factor was negative and non-significant. In addition, many of the factor loadings’ standard errors were much larger than in the four correlated factors model.

**Table 5.** Standardized factor loadings and factor correlations from Model 5h and its corresponding bifactor model

| Disorder/Trait | Four Correlated Factors Model (Model 5h) |  |  |  |  | Bifactor Model with Four Specific Factors (Model 5i) |  |  |  |  |
| --- | --- | --- | --- | --- | --- | --- | --- | --- | --- | --- |
|  | EXT | INT | TP | NDD |  | GEN | EXT | INT | TP | NDD |
| Antisocial Behavior | .64 (.14)*** |  |  |  |  | .55 (.10)*** | .29 (.75) |  |  |  |
| Tobacco | .43 (.09)*** |  |  |  |  | .34 (.07)*** | .15 (.40) |  |  |  |
| Alcohol | .75 (.13)*** |  |  |  |  | .60 (.06)*** | .80 (2.01) |  |  |  |
| Cannabis | .47 (.16)** |  |  |  |  | .41 (.13)** | -.10 (.43) |  |  |  |
| PTSD |  | .56 (.09)*** |  |  |  | .68 (.15)*** |  | -.69 (.68) |  |  |
| MDD |  | .95 (.10)*** |  |  |  | .94 (.08)*** |  | .31 (.25) |  |  |
| Anxiety |  | .72 (.09)*** |  |  |  | .67 (.12)*** |  | .50 (.33) |  |  |
| Eating Disorder |  |  | .25 (.05)*** |  |  | -.05 (.06) |  |  | .32 (.06)*** |  |
| Schizophrenia |  |  | .90 (.05)*** |  |  | .36 (.03)*** |  |  | .78 (.08)*** |  |
| Bipolar Disorder |  |  | .76 (.04)*** |  |  | .30 (.03)*** |  |  | .73 (.08)*** |  |
| OCD |  |  | .39 (.05)*** |  |  | .15 (.07)* |  |  | .38 (.07)*** |  |
| ASD |  |  |  | .52 (.05)*** |  | .41 (.05)*** |  |  |  | .29 (.12)* |
| Aggression |  |  |  | .74 (.12)*** |  | .44 (.12)*** |  |  |  | .86 (.36)* |
| ADHD |  |  |  | .77 (.06)*** |  | .54 (.05)*** |  |  |  | .50 (.21)* |
| EXT | — |  |  |  |  | — | — | — | — | — |
| INT | .66 (.14)*** | — |  |  |  | — | — | — | — | — |
| TP | .41 (.08)*** | .42 (.05)*** | — |  |  | — | — | — | — | — |
| NDD | .67 (.15)*** | .75 (.09)*** | .19 (.04)*** | — |  | — | — | — | — | — |

**Note.** ADHD = attention/deficit-hyperactivity disorder, ASD = autism spectrum disorder, MDD = major depressive disorder, PTSD = post-traumatic stress disorders, OCD = obsessive compulsive disorder, EXT = Externalizing factor, INT = Internalizing factor, TP = Thought Problems factor, NDD = Neurodevelopmental Disorders factor, GEN = General factor, $\beta$ = Standardized regression coefficient, SE = Standard Error. \* $p < .05$ , \*\* $p < .01$ , \*\*\* $p < .001$ .

Our results thus suggest that the best-fitting model comprises four moderately correlated factors of externalizing, internalizing, thought problems, and neurodevelopmental disorders, in which eating disorders loads only on thought problems (Model 5h). As shown in Figure 1 and Table 5, all factor loadings and correlations were significant, as they were greater than twice their standard errors, and were moderate to high. The exception to this was eating disorders, which had a small but significant loading on thought problems. The average of the disorders’ and traits’ genetic variance accounted for by the factors was substantial (internalizing = .54, externalizing = .33, thought problems = .38, and neurodevelopmental disorders = .49).

**Figure 1.**
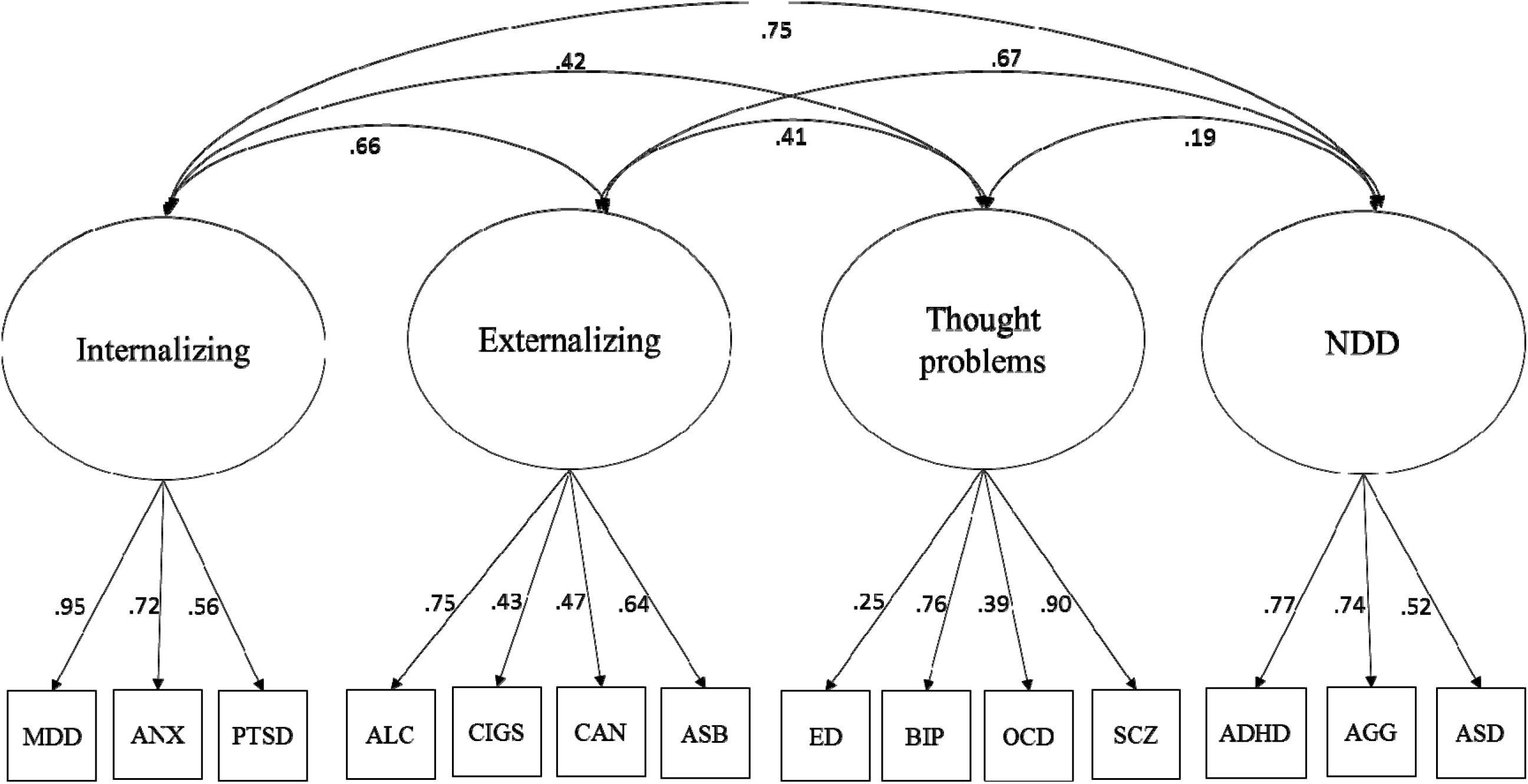
Best-fitting confirmatory factor analysis model. NDD – neurodevelopmental disorders, SCZ – schizophrenia, OCD – obsessive-compulsive disorder, BIP – bipolar disorder, ASD – autism spectrum disorder, PTSD – post-traumatic stress disorder, MDD – major depressive disorder, ANX – anxiety disorders, ED – eating disorders, ASB – antisocial behavior, ALC – alcohol dependence, CAN – cannabis dependence, CIGS – number of cigarettes smoked per day, ADHD – attention-deficit/hyperactivity disorder, AGG – aggression

The externalizing and internalizing factors were positively and moderately correlated with all other factors and with each other, while thought problems and neurodevelopmental disorders were only weakly correlated. As shown in Figure 1, the neurodevelopmental disorders factor was moderately to highly genetically correlated with the externalizing and internalizing factors (.67 and .75, respectively), suggesting that the genes that predispose to neurodevelopmental disorders in early childhood also predispose to externalizing and internalizing disorders later in childhood and into adolescence and adulthood.

Figure 2 presents the differential genetic correlations between the higher-order psychopathology dimensions from Model 5h and the external criterion variables listed in Table 2. The externalizing factor was more strongly correlated with extraversion, age at first birth (negative), and educational attainment (negative) than were the internalizing and neurodevelopmental disorders factors. The thought problems dimension had the weakest correlations with these external variables. The externalizing and neurodevelopmental disorders dimensions were more strongly correlated with total number of children born than were the internalizing or thought problems dimensions.

**Figure 2.**
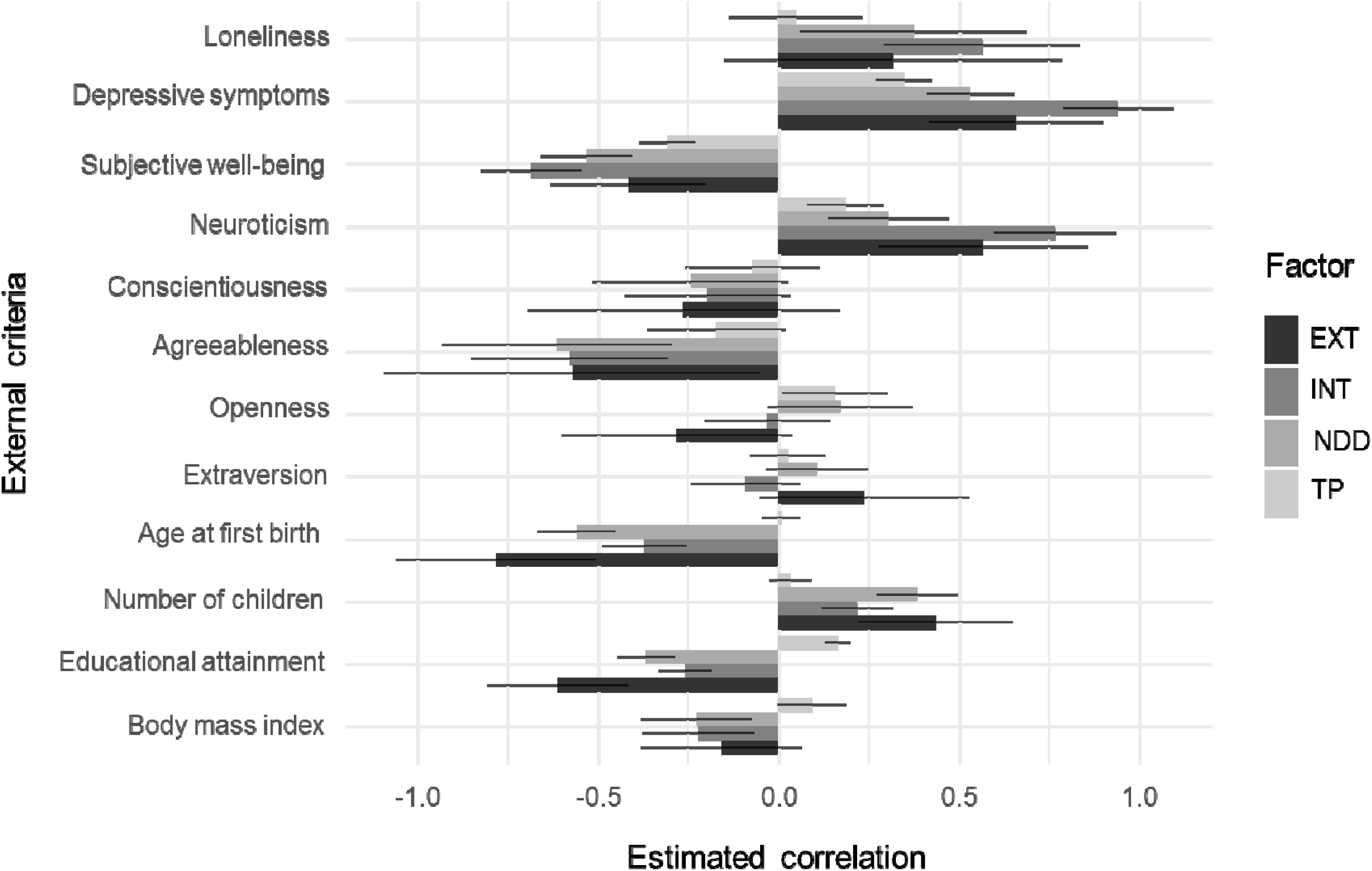
Genetic correlations of the external criterion variables with the four higher-order psychopathology factors. EXT – externalizing higher-order factor, INT – internalizing higher-order factor, TP – thought problems higher-order factor, NDD – neurodevelopmental disorders higher-order factor. Bars indicate 95% confidence intervals

In contrast, the internalizing factor was strongly related to loneliness, depression symptoms, subjective well-being (negative), and neuroticism. The externalizing and neurodevelopmental disorders factors were more strongly associated with these criteria than thought problems. Externalizing, internalizing, and neurodevelopmental disorders had similar negative associations with conscientiousness, agreeableness, and body mass index. Finally, thought problems was positively correlated with openness to experience and educational attainment, whereas the other factors were either unrelated or negatively related.

Most crucially, the direction of associations between the four higher-order psychopathology factors and the external criteria were in the expected direction, and the relative magnitude of the four factors’ genetic correlations with the external criteria also matched theoretical expectations. For example, all psychopathology dimensions had some association with loneliness, depressive symptoms, and subjective well-being, but the internalizing factor displayed the largest associations. These different patterns of genetic correlations provide evidence for the external validity of the higher-order psychopathology factors.

## DISCUSSION

In this study, factor analyses of GWAS summary statistics for fourteen psychiatric disorders and related traits revealed four moderately correlated factors – externalizing, internalizing, thought problems, and neurodevelopmental disorders – which showed distinct patterns of genetic correlations with external validity variables. A bifactor model comprising a general factor of psychopathology did not fit as well as the corresponding best-fitting correlated factors model and yielded problematic model characteristics, suggestive of overfitting.

Given that our analyses used GWAS summary statistics from fourteen different studies, it is noteworthy that our best-fitting model mirrored features found in many phenotypic factor analyses, such as moderate-to-high factor loadings and moderate factor correlations. Indeed, these features and the four factors in our best-fitting model mirror crucial aspects of the HiTOP model of psychopathology.

In addition, each GWAS comprises meta-analyses of distinct cohorts, rather than a single cohort in which participants reported on all disorders simultaneously. Our analyses are thus unaffected by issues such as shared measurement error, response biases, common method variance, and small sample size that can affect phenotypic studies.

It is worth noting that many of the models we tested fell short of conventional standards for good model fit, likely due to the limitations of extant GWAS summary statistics. However, the best fitting model did surpass standards for good fit (i.e., CFI > 0.90). The four factors were also differentially associated with external criterion variables, suggesting that they represent meaningfully distinct dimensions of psychopathology. The direction and magnitude of these correlations is consistent with previous phenotypic^45,46^ and genetic studies^31,47–49^ of higher-order dimensions of psychopathology.

Finally, the neurodevelopmental disorders factor was moderately-to-highly genetically correlated with the externalizing and internalizing factors, suggesting that the genes that predispose to neurodevelopmental disorders in early childhood also predispose to externalizing and internalizing disorders later in childhood and into adolescence and adulthood. This suggests an etiological basis for the association of ADHD or ASD in childhood with antisocial behavior, substance use disorders, anxiety, and depression in adolescence and adulthood.

Two previous studies have modeled the structure of psychopathology using GWAS summary statistics^19,20^, and our results strengthen the main conclusion of those studies that factor analysis can be used to model genetic covariation among psychopathological disorders. The current study adds to this literature by including a greater number of psychiatric disorders and related psychological traits in the analyses and testing a greater number of alternative models of psychopathology.

The best-fitting model in the current study indicated that eating disorders and OCD, on the one hand, and bipolar disorder and schizophrenia, on the other, load together on the same thought problems factor, which mirrors findings from a previous study^19^. We also replicated the finding that ASD and ADHD load together on a separate neurodevelopmental disorders factor^20^.

Nevertheless, our results differ from these two previous studies in a number of important ways. First, using CFA, we found that the fourteen disorders and related traits included in our study were best represented by four correlated factors, including a thought problems factor onto which bipolar disorder, schizophrenia, OCD, and eating disorders loaded. Second, our findings suggest that MDD loads together with other internalizing disorders rather than with bipolar disorder and schizophrenia. These differences across studies illustrate how the inclusion or exclusion of particular disorders or traits, as well as the use of different statistical methods, can yield different results.

For disorders whose placement in the multivariate higher-order structure of psychopathology is still open to debate, we tested alternative models in which the disorder loaded on multiple factors. The most notable such modification is the placement of eating disorders, which ultimately loaded on the thought problems factor. Recently, structural models of psychopathology suggested that eating disorders can be placed within the internalizing framework^50,51^, although some models suggest it is a separate dimension^8^. Our finding that these disorders loaded most strongly on the thought problems factor seems to suggest that this factor is characterized by disturbed cognitions found across disparate psychopathological disorders. The placement of eating disorders on this factor is consistent with previous studies that have found substantial covariation between eating disorders and OCD^52,53^, which also loaded on the thought problems factor in the current study.

Our findings can also be contextualized within the current literature on the phenotypic structure of psychopathology. The HiTOP model includes most forms of psychopathology, several of which have not been studied in a GWAS and were thus not included in the current analyses. However, comparison of our results with the HiTOP model yields some interesting points. First, the HiTOP model, and indeed other phenotypic models of psychopathology^54,55^, distinguish between disinhibited (e.g., substance use) and antagonistic (e.g., antisocial personality and other personality disorders) forms of externalizing. In our analyses, however, a model that distinguished substance use pathology from other externalizing disorders did not perform well.

Second, in the HiTOP framework, eating disorders and OCD are clustered within internalizing psychopathology, whereas they were best characterized within the thought problems factor in the current study. Finally, the HiTOP model tentatively posits that mania can be captured within both internalizing and thought disorder factors. Our inability to distinguish between mania and depression within bipolar disorder precluded a test of this model. Rather, bipolar disorder loaded with other thought disorders, perhaps reflecting the strong genetic relationship between more severe mania and schizophrenia^3^.

As GWAS summary statistics become available on more fine-grained dimensions of psychopathology, we will be able to test more detailed models posited within the HiTOP framework, such as distinguishing between fears and distress pathology within internalizing, and modeling dimensions of detachment and somatoform psychopathology.

Modeling higher-order psychopathology dimensions may have several advantages for genetic studies over studying individual diagnoses one at a time. These include a more parsimonious and accurate representation of psychopathology^8,56^, higher heritability, capitalization on pleiotropy to increase genetic associations^57,58^, greater genetic correlations with external variables, greater statistical power to detect genetic associations due to more information contained in latent continuous versus observed categorical phenotypes^54,55^, and elimination of measurement error.

These advantages, as well as GWAS of more fine-grained phenotypes (e.g., of distinct anxiety disorders), should increase the genetic signal and consequently the number of genome-wide significant associations found in GWAS^59^. The resolving power of such factor analyses should increase as individual GWAS meta-analyses become larger and better powered statistically, and as continuous psychopathology dimensions are included as phenotypes in GWAS, both of which should result in higher GWAS-based heritabilities and greater genetic signal to model.

Future research using GWAS of continuous psychopathology dimensions in large samples should attempt to replicate the higher-order structure of psychopathology presented in this study.

## ACKNOWLEDGEMENTS

Earlier versions of this work were presented at the 32nd Meeting of the Society for Research in Psychopathology and the 49th Meeting of the Behavior Genetics Association. I.D. Waldman, H.E. Poore and J.M. Luningham contributed equally to this work.

